# Classifying high-dimensional phenotypes with ensemble learning

**DOI:** 10.1101/2023.05.29.542750

**Authors:** Jay Devine, Helen K. Kurki, Jonathan R. Epp, Paula N. Gonzalez, Peter Claes, Benedikt Hallgrímsson

## Abstract

1. Classification is a fundamental task in biology used to assign members to a class. While linear discriminant functions have long been effective, advances in phenotypic data collection are yielding increasingly high-dimensional datasets with more classes, unequal class covariances, and non-linear distributions. Numerous studies have deployed machine learning techniques to classify such distributions, but they are often restricted to a particular organism, a limited set of algorithms, and/or a specific classification task. In addition, the utility of ensemble learning or the strategic combination of models has not been fully explored.
2. We performed a meta-analysis of 33 algorithms across 20 datasets containing over 20,000 high-dimensional shape phenotypes using an ensemble learning framework. Both binary (e.g., sex, environment) and multi-class (e.g., species, genotype, population) classification tasks were considered. The ensemble workflow contains functions for preprocessing, training individual learners and ensembles, and model evaluation. We evaluated algorithm performance within and among datasets. Furthermore, we quantified the extent to which various dataset and phenotypic properties impact performance.
3. We found that discriminant analysis variants and neural networks were the most accurate base learners on average. However, their performance varied substantially between datasets. Ensemble models achieved the highest performance on average, both within and among datasets, increasing average accuracy by up to 3% over the top base learner. Higher class R^2^ values, mean class shape distances, and between– vs. within-class variances were positively associated with performance, whereas higher class covariance distances were negatively associated. Class balance and total sample size were not predictive.
4. Learning-based classification is a complex task driven by many hyperparameters. We demonstrate that selecting and optimizing an algorithm based on the results of another study is a flawed strategy. Ensemble models instead offer a flexible approach that is data agnostic and exceptionally accurate. By assessing the impact of various dataset and phenotypic properties on classification performance, we also offer potential explanations for variation in performance. Researchers interested in maximizing performance stand to benefit from the simplicity and effectiveness of our approach made accessible via the R package *pheble*.

## 1.0 Introduction

Linear discrimination methods have long been used in quantitative phenotypic analyses to visualize and discriminate classes. Linear discriminators (e.g., linear discriminant analysis) tend to be efficient and sufficiently accurate on low-dimensional datasets, such as those with a few linear measurements or small, sparse landmark configurations (Mitteroecker & Bookstein, 2011). However, advances in data collection techniques (Devine et al., 2020; Percival et al., 2019; Porto et al., 2021) and data crowdsourcing (Boyer et al., 2016) are yielding increasingly large, high-dimensional phenotypic datasets with more classes, unequal class covariances, and non-linear distributions. Non-parametric machine learning approaches have been developed to classify such distributions, and numerous self-contained studies have hinted at their potential (Lürig et al., 2021), but the utility of these methods for classifying high-dimensional phenotypes has not been systematically investigated on a large scale. Because traditional machine learning models often fail to achieve satisfactory performance when dealing with certain data structures (e.g., noisy, imbalanced, etc.), it is further worth considering how ensemble learning or the strategic integration of these models can improve performance. In this paper, we present a comprehensive analysis of learning-based classification algorithms on a collection of morphometric datasets and show how ensemble learning can maximize discrimination in arbitrary biological settings.

Classification is the process of assigning members to a class. This task can be accomplished through different learning strategies. Ensemble learning, and blending in particular, is our focus. Blending ensemble approaches involve strategically stacking a set of individual classifiers using a holdout validation set to improve performance (Breiman, 1996; van der Laan et al., 2007). Each classifier alone is relatively simple and easy to train, often only performing well on a subset of the data, but together these weak classifiers become a strong classifier. Despite its success in other fields, ensemble learning has rarely been explored in phenomics due to the paucity of open-source implementations, insufficient expertise, and a continual reliance on the same methods. For example, linear discriminant analysis, the hallmark approach to phenotypic classification, maximizes the ratio of between-class variance to within-class variance to ensure maximal separability. Unfortunately, this method assumes equality of covariances among classes and can only find a linear discriminant function (i.e., a linear combination) to separate them (Mitteroecker & Bookstein, 2011; Sheets et al., 2006). While homoscedasticity is common among datasets with only a few groups, larger phenotypic datasets with heterogeneous groups stand to benefit from non-parametric alternatives.

Recent applications of learning-enabled classification for high-dimensional phenotypes have either involved a single dataset (e.g., one species or one study) (Hosseini et al., 2019; Salifu et al., 2022), small sample sizes (Courtenay et al., 2019; Courtenay and González-Aguilera, 2020), a specific learning problem (e.g., only binary or multi-class classification with a single dataset) (Courtenay et al., 2019, 2021), and/or a single algorithm (Bertsatos et al., 2020; Fellowes et al., 2019). As such, there has not been a detailed examination of these machine learning algorithms under different biological conditions. There have also been few attempts at combining multiple base learners into a strong phenotypic learner via blending or stacking, a similar technique in ensemble learning. The *H2O* (Candel et al., 2016), *SuperLearner* (Polley et al., 2019), and *caretEnsemble* (Deane-Mayer & Knowles, 2016) R packages offer tools for ensemble learning, but they lack either (a) a large, diverse library of classification algorithms, (b) multi-class ensemble capabilities, and/or (c) a streamlined ensemble workflow for non-experts. Rather than conduct one-off studies, it is important to test learning-based methods with diverse high-dimensional phenotypic datasets and a standardized workflow.

We present an empirical analysis of 33 learning-based classification algorithms and various blending ensembles across 20 high-dimensional morphometric datasets using a new R package, *pheble*. We examine a variety of algorithm families, including Bayesian methods, decision trees, bagging and boosting ensembles, kernel-based methods, neural networks, and regression methods. Binary and multi-class classification tasks central to evolutionary biology, developmental biology, and ecology are considered. Specifically, we attempt to discriminate sex and different environmental classes in the binary classification experiments, then turn to classes such as species, population, genotype, and habitat in the multi-class experiments. To investigate potential determinants of classification accuracy, including class R^2^ values, unequal class covariances, mean class shape distances, between– vs. within-class class variances, class imbalances, and sample size, we employ phenotypic datasets containing a range of anatomical data from different organisms with unique class distributions. Ultimately, we illustrate how ensemble models outperform all other base learners on average whilst being consistently accurate. Our code is freely available at github.com/jaydevine/pheble.

## 2.0 Materials and Methods

### 2.1 Datasets

We use 20 publicly available morphometric datasets to complete a classification meta-analysis and test the viability of an ensemble workflow. Table 1 enumerates the key metadata. Additional information about data provenance is listed in Table S1. Altogether these datasets represent a wide assortment of families, ranging from small, terrestrial insects (e.g., *Formicidae*) to large, aquatic mammals (e.g., *Crocodylidae*) with distinct anatomies, class distributions, and sample sizes.

**Table 1.** Summary of phenotypic datasets, including the family (i.e., dataset name), landmarked anatomy, total sample size (*N*), class, number of class levels, and number of phenotypic variables. The “/” delimiter indicates datasets with two families, whereas the “+” suffix indicates datasets with more than three families.

| Family | Anatomy | $N$ | Class | Levels | Variables |
| --- | --- | --- | --- | --- | --- |
| *Asterinidae* | Body | 885 | Sex | 2 | 20 |
| *Drosophilidae* | Wing | 2926 | Sex | 2 | 96 |
| *Emydidae* | Shell | 2161 | Habitat | 2 | 159 |
| *Gasterosteidae* | Skull | 190 | Habitat | 2 | 210 |
| *Gasterosteidae* | Body | 521 | Sex | 2 | 30 |
| *Hominidae* | Sacrum | 101 | Sex | 2 | 300 |
| *Hynobiidae/Cryptobranchidae* | Palate | 62 | Habitat | 2 | 48 |
| *Muridae* | Cranium | 1251 | Sex | 2 | 2532 |
| *Poeciliidae* | Body | 1449 | Sex | 2 | 26 |
| *Serranidae/Sparidae* | Body | 259 | Site | 2 | 26 |
| *Cichlidae* | Jaw | 1136 | Tribe | 14 | 126 |
| *Colubridae*+ | Vertebrae | 1260 | Species | 15 | 24 |
| *Crocodylidae/Alligatoridae* | Cranium | 183 | Species | 8 | 234 |
| *Drosophilidae* | Wing | 2926 | Elevation | 9 | 96 |
| *Formicidae* | Face | 1494 | Species | 6 | 22 |
| *Muridae* | Cranium | 1251 | Genotype | 26 | 2532 |
| *Ocypodidae* | Carapace | 1867 | Species | 16 | 42 |
| *Percidae* | Body | 423 | Species | 15 | 20 |
| *Vespidae* | Wing | 206 | Species | 8 | 38 |
| *Viviparidae* | Shell | 1224 | Population | 22 | 254 |

For binary classification, we mainly concentrate on sex discrimination, but other classes (e.g., habitat or site) are incorporated to experiment with different classifiers. Likewise, we primarily focus on species discrimination for multi-class classification, but additional classes (e.g., population, genotype, or habitat/elevation) are included for experimentation. Each dataset is composed of a sparse or dense array of *p* homologous anatomical landmarks in *k* dimensions, resulting in *p* × *k* phenotypic variables for every observation (Table S1). Using the *Morpho* (Schlager, 2017) and *geomorph* (Adams & Otárola-Castillo, 2013) R packages, we superimpose the landmark configurations into a common shape space for each dataset via Generalized Procrustes Analysis (GPA) (Gower, 1975; Rohlf & Slice, 1990) to obtain Procrustes shape coordinates.

### 2.2 The R package pheble

The R package *pheble* contains functions to build a streamlined ensemble learning workflow for classifying high-dimensional data (Fig. 1). Typically, this involves (1) preprocessing a dataset, (2) training a multitude of models to perform a given classification task, (3) strategically selecting and combining those model predictions to train an ensemble model, and (4) evaluating the models on an unseen dataset. We describe each step in detail below.

**Figure 1.**
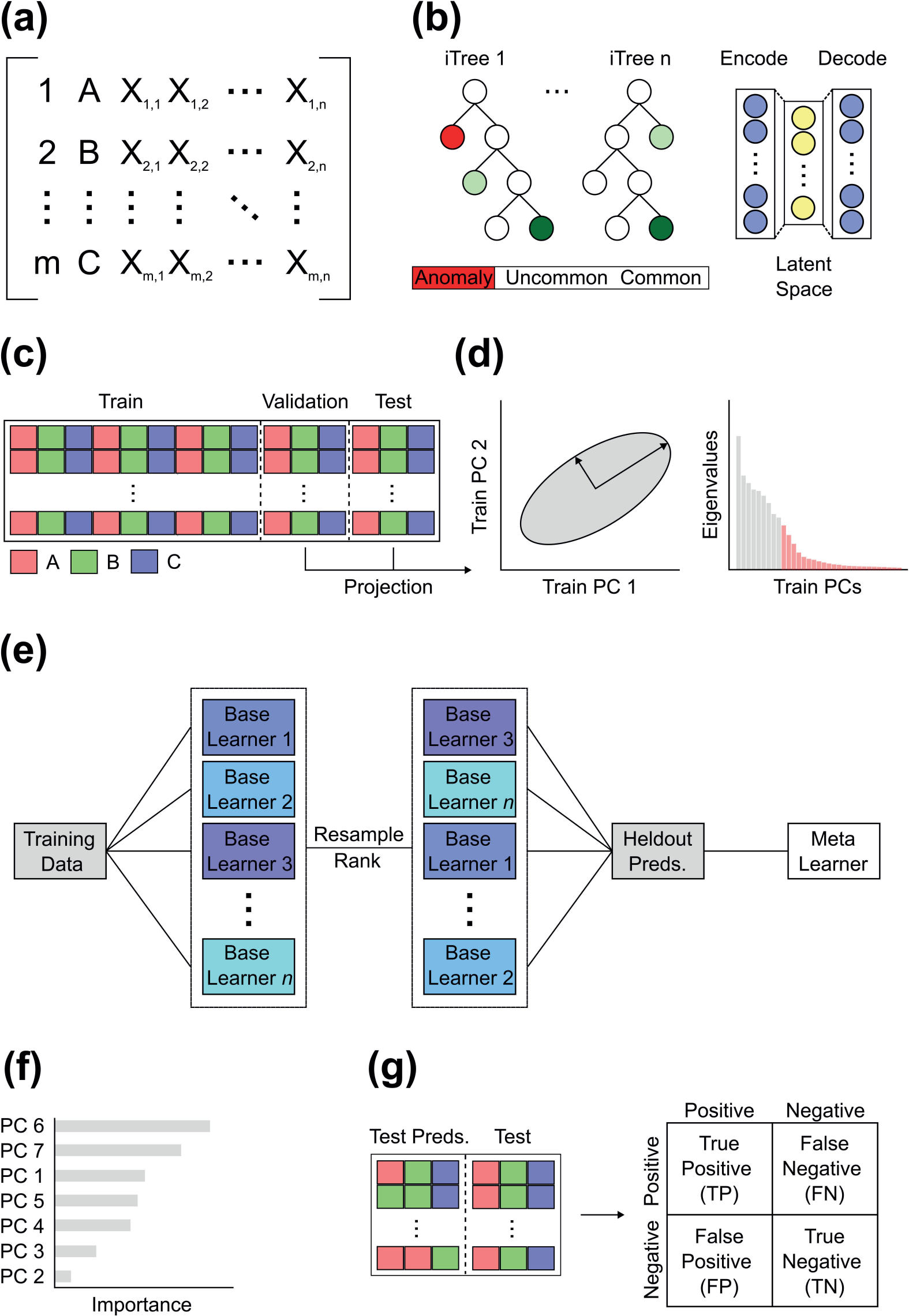
Schematic overview of ensemble learning workflow. (a) A matrix containing observation names, classes, and high-dimensional data is (b) preprocessed with an (left) extended isolation forest or (right) autoencoder to remove anomalies, then (c) split into training, holdout validation, and test sets, ensuring proportional class representations. (d) Dimensionality reduction is performed on the training set and higher order principal components are removed according to a variance threshold before projecting the validation and test data into that space. (e) An arbitrary number of base learners are trained on the training data, ranked according to the resampling optimization metric, and the top x learners are applied to the holdout set to generate predictions. The metalearner is trained on these predictions and later used to predict the test classes. (f) A variable importance readout is returned, along with (g) evaluation results for every method via the confusion matrix.

### 2.3 Preprocessing

#### 2.3.1 Anomaly detection

Anomaly detection is the process of finding patterns in the data that do not conform to expected behavior (Chandola et al., 2009). We provide autoencoder and extended isolation forest options for anomaly detection using algorithms in the *H2O* R package, as these methods are effective and generic enough to handle most data (Fig. 1b). Autoencoders are a type of neural network designed to encode the input data into compressed but meaningful representations, often called latent variables, then decode them back into a reconstructed output that is as similar as possible to the input (Hinton & Salakhutdinov, 2006). Poor reconstructions have higher errors and are indicative of anomalies. Isolation forests, by contrast, utilize a tree structure with branches built from random cuts or thresholds in the values of randomly selected features (Liu et al., 2012). Because the branching process can introduce bias, the extended variant was proposed (Hariri et al., 2019). The deeper a sample travels into these branches, the less likely it is to be anomalous.

We implement random discrete grid searches to optimize the hyperparameters of both anomaly detection methods. This involves iteratively testing random combinations of hyperparameters to a user-defined tune length, set here to 100, then evaluating each model and selecting the one with the lowest mean squared error. We evaluate the efficacy of each approach by correlating their anomaly scores with Procrustes distances to the mean, the most widely accepted measure for outlier detection in morphometrics. The average autoencoder correlation is *r* = 0.85 (Fig. S1), whereas the average extended isolation forest correlation is *r* = 0.82 (Fig. S2). Despite these promising results, we stick to the Procrustes convention and only remove anomalies based on the Procrustes distance interquartile range. Users should feel confident generalizing these methods to non-morphometric data when a straightforward measure is unavailable.

#### 2.3.2 Data partitioning and dimensionality reduction

The training set enables a model to learn underlying patterns and relationships in the data, while the test set facilitates unbiased evaluations of a final model. We invoke a type of ensemble learning called blending, where a validation set is partitioned from the training set to generate an initial collection of predictions to train a metalearner (Fig. 1c). By combining held-out predictions from multiple, usually diverse, base learners, the metalearner develops into a single, secondary prediction model with more discriminative power (LeDell, 2015). The metalearner is not limited to any particular algorithm, although generalized linear models and random forests to a lesser extent tend to be employed, as they are more resistant to overfitting (LeDell et al., 2016). We apply a 70/15/15% training, validation, and test split to each dataset in this study. We ensure that class levels are represented sufficiently and proportionately across the partitioned datasets (Fig. 1c).

High-dimensional data are information rich but tend to be redundant or highly correlated, leading to inefficient models with noise and less discriminative power. To decompose these datasets, we provide Principal Component Analysis (PCA) and autoencoder dimensionality reduction options in the data partitioning function (Fig. 1d); however, other extracted features or even the raw data can be defined as inputs. While PCA continues to reign supreme when studying between-class differences via linear decomposition (Du, 2019), phenotypic traits can exhibit non-linear relationships (e.g., Unger et al., 2021), in which case an autoencoder might be preferable. After the data are partitioned, dimensionality reduction is performed on the training set, then the validation and test sets are predicted with that model (Figs. 1c,d). We use PC scores as training, validation, and test data due to the highly correlated and Euclidean nature of Procrustes coordinates projected into tangent space.

### 2.4 Training

Ensemble models benefit from a comprehensive library of base learners (Fig. 1e). Since existing R packages lack either multi-class ensemble capabilities or a large enough selection of base learners, we leverage training algorithms from *caret* (Kuhn, 2008), the most celebrated and comprehensive machine learning classification package in R. After experimenting with every parametric and non-parametric supervised learning method, we homed in on 33 learners for binary classification and 30 learners for multi-class classification. Algorithms with excessive training times and susceptibility to errors were excluded. Table 2 lists the major algorithm families and associated algorithms. Detailed information about these algorithm families can be found in various reviews (Mitteroecker & Bookstein, 2011; Lürig et al., 2021).

**Table 2.** Summary of algorithm families and available algorithms. Acronym denotes the name in R. Each algorithm is capable of binary and multi-class classification, except those with an asterisk (*), which are unavailable for multi-class tasks.

| Algorithm family | Algorithm and acronym |
| --- | --- |
| Bayesian | Naïve Bayes (nb) |
| Decision trees | C5.0, conditional inference forest (cforest), evolutionary decision tree (evtree), random forest (rf or ranger) |
| Ensemble (bagging/boosting) | AdaBoost.M1, AdaBag, multivariate adaptive regression spline with bagging and boosting (bagEarthGCV), Classification and Regression Tree with bagging (treebag) |
| Kernel/instance-based | Discriminant analysis (flexible (fda), heteroscedastic (hda), high-dimensional (hdda), linear (lda), localized (loclda), mixture (mda), penalized (pda), quadratic (qda), regularized (rda), stepwise linear (stepLDA), stepwise quadratic (stepQDA), sparse linear (sparseLDA)), Gaussian process (linear (*gaussprLinear), polynomial (*gaussprPoly), radial (*gaussprRadial)), *k*-nearest neighbors (kknn), support vector machine (linear (svmLinear), polynomial (svmPoly), radial (svmRadial)) |
| Neural networks | Artificial neural network (nnet) |
| Regression | Generalized linear model (glmnet), multivariate adaptive regression spline (earth), partial least squares (pls) |

Before training a model, it is wise to introduce a resampling strategy. Oftentimes, a list of training models will be evaluated on the training data prior to testing. But evaluating a model on the full training dataset inflates the initial evaluation metrics and offers no insight into model generalization on new datasets. Much like bagging, it is more instructive to repeatedly resample the training data (e.g., via bootstrapping or cross-validation), train on the subsample, predict on the held-out sample, and average the results to arrive at a representative understanding of model performance. This is a general feature of *caret* that we integrate. Optionally, the resampling can include an up– or down-sampling step to redress class imbalances. We employ a bootstrapping (N=25 iterations) option across each iteration of the hyperparameter optimization, but also make cross-validation variants available.

A well-trained model is heavily dependent on hyperparameter optimization. While manually defining and optimizing a list of custom hyperparameters is feasible for a single learner, it is tedious and time-consuming to do the same for myriad learners. Again, we capitalize on the automatic hyperparameter tuning capabilities of *caret* and allow users to specify the tune length of each base learner. We set the tune length to 10 and perform a random discrete search, meaning a maximum of 10 random hyperparameter combinations are evaluated for each model. The best model according to a user-defined metric is retained. We define ROC and Cohen’s Kappa as the default metrics for binary and multi-class classification, respectively. However, log loss, accuracy, balanced accuracy, and F1 metrics are additionally available.

Higher resampling iterations and tune lengths will not only increase the generalizability of a model but also the likelihood of reaching an optimum. Unfortunately, training time rises exponentially if these values are set too high, because they are applied to every base learner. If the task is not time-sensitive, doubling or even tripling the resampling and tune length numbers should be feasible. Under time constraints, however, the values proposed above should be sufficient, though they can certainly be decreased if the dataset and/or parameter space is massive. Fig. S3 shows the distribution of training times for each algorithm. To accelerate training, we provide an argument to specify the number of cores for parallelization. Invoking as many cores as possible will dramatically reduce training times. We trained every model using 10 cores on an Intel i7– 8700K Processor (3.70 GHz).

After compiling a list of successfully trained models, we rank order them using the optimization metric from the resampling process (Fig. 1e). Anywhere between two and the total number of successful base learners can be selected for the ensemble. Models that do not converge or fail to predict complete cases are eliminated. For computational reasons, we choose the top three, top five, and top 10 models to construct multiple ensembles. Each top learner is initially deployed to predict the classes of the validation and test sets, then the held-out validation predictions are stacked to train each ensemble. We train the ensembles according to the procedures above, except with a generalized linear model or random forest metalearner. We prioritize these metalearners for their robustness to overfitting, but any algorithm can be used. Equipped with the test set predictions as new test data, we predict the test classes with the ensemble. We additionally predict the test classes from each successful base learner after feeding them the original test data to provide a comparative summary of method performance. Overlap between the predicted and observed test classes is evaluated using confusion matrices (Fig. 1g).

### 2.5 Variable importance

Predictors tend to vary in their ability to discriminate classes. Explainability or understanding the relative importance of each variable to a model is thus helpful, particularly in high-dimensional space where teasing apart effects is difficult. Quantifying importances from a single classification model is easily accomplished with existing functions. However, there is no standard approach for re-weighting them in an ensemble. We therefore compute and store the original variable (e.g., PC) importances from every individual base learner in the ensemble, multiply these importances by the corresponding model importances from the held-out validation predictions, then calculate the weighted mean importance of each variable (Fig. 1f).

### 2.6 Evaluation metrics

We acquire a standard set of classification metrics for each base learner and ensemble using the confusion matrix. While the metrics below primarily concern the test data, we also gather the same metrics for the validation data to understand the composition of the ensemble. Hereafter, we focus on F1 scores and balanced accuracy, as they measure overall model performance by incorporating precision, recall, sensitivity, and specificity. But other measures, including positive prediction value, negative prediction value, prevalence, detection rate, detection prevalence, accuracy, and Kappa, are provided as a function output. Both F1 and balanced accuracy can be expressed as ratios between the number of true positives (TP), true negatives (TN), false positives (FP), and false negatives (FN) (Fig. 1g):

*precision* = *TP* / (*TP* + *FP*)

*sensitivity* = *TP* / (*TP* + *FN*)

*specificity* = *TN* / (*TN* + *FP*)

*F*1 = 2 · (*precision* · *sensitivity*) / (*precision* + *sensitivity*)

*balanced accuracy* = (*sensitivity* · *specificity*) / 2

F1 emphasizes the number of true positives or correctly predicted positive classes relative to the total number of predictions, whereas balanced accuracy accounts for both true positives and true negatives. We examine variation in F1 scores and balanced accuracy among datasets (i.e., within methods) and within datasets (i.e., among methods) after separating the binary and multi-class classification results. We then interrogate possible causes of performance variation by merging the classification results. With F1 or balanced accuracy as the response variable and classification task (binary/multi) plus class R^2^, mean class covariance distance, mean class shape distance, between– vs. within-class variance, class balance, or sample size as the explanatory variables, we fit multiple regression models. Class R^2^ is the R^2^ value obtained from fitting a linear model, with residual randomization, of Procrustes shape coordinates on class (Collyer & Adams, 2018). Mean class covariance distance is the mean Euclidean distance between the covariance matrices of every unique pairwise class combination (Le Maître & Mitteroecker, 2019). Mean class shape distance is the mean Procrustes distance between the mean shapes of every unique pairwise class combination. Between– vs. within-class variance is the quotient of the traces of the between– and within-class covariance matrices (Le Maître & Mitteroecker, 2019). Class balance is a summary measure for the number of observations per class relative to the sample size and is measured as the Shannon entropy normalized by the number of classes. Sample size is the total sample size.

## 3.0 Results

We classified nearly 10,000 high-dimensional shape phenotypes from 10 binary class datasets using 33 classification algorithms and their ensembles. Fig. 2a shows the distribution of F1 scores and balanced accuracies for the top 10 base learners and select ensembles among binary class datasets (see Table S2 and Fig. S4 for all algorithms). For base learners, the top 10 average F1 scores in descending order were attained by regularized discriminant analysis (rda), heteroscedastic discriminant analysis (hda), neural network (nnet), localized linear discriminant analysis (loclda), AdaBoost, sparse linear discriminant analysis (sparseLDA), partial least squares (pls), mixture discriminant analysis (mda), quadratic discriminant analysis (qda), and polynomial support vector machine (svmPoly). The top three and top five random forest (rf) ensembles ranked first and third among all methods, averaging 91.4% and 90.7% F1 scores, respectively, in between which rda achieved 91.0%. While the top 10 generalized linear model (glm) ensemble ranked fourth at 90.5%, the top three and top five glm ensembles dropped to 84.8% and 84.7%, respectively. As for average balanced accuracy, the top 10 base learners were rda, hda, loclda, svmPoly, mda, nnet, radial support vector machine (svmRadial), AdaBoost, sparseLDA, and pls.

**Figure 2.**
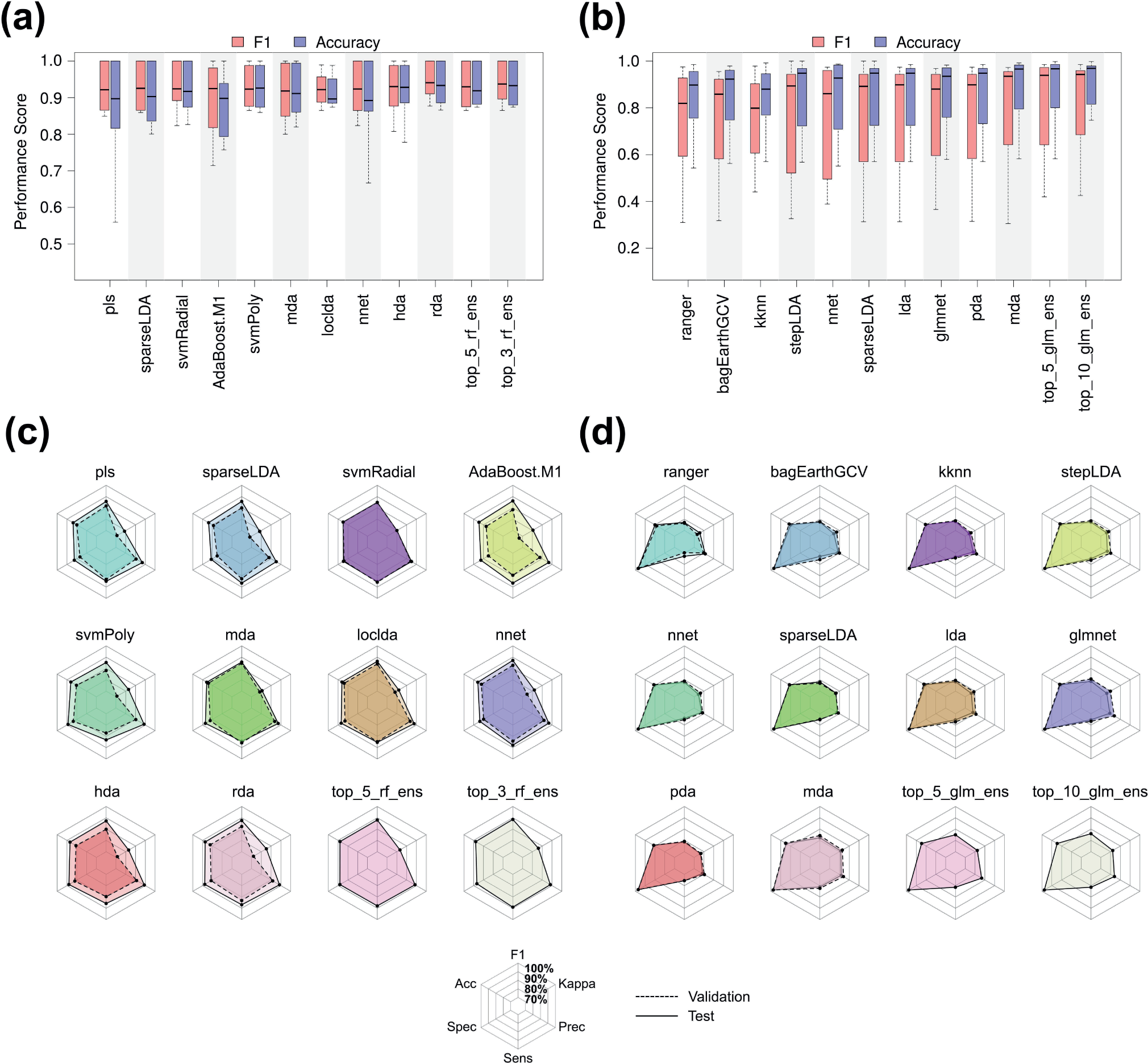
(a) Binary and (b) multi-class F1 (red) and balanced accuracy (blue) distributions for the top 10 base learners and ensembles, shown in ascending order. Other ensembles are excluded for simplicity. (c) Binary and (d) multi-class radar plots detailing average performance for the validation (dashed line) and test (solid line) set data across the same base learners and select ensembles, shown in ascending order. Radar plot lines start at 70% and radiate outward towards 100%.

The top three and top five rf ensembles ranked second and third with 90.9% and 90.5% balanced accuracies, respectively, behind the leading 91.0% of rda.

We also classified roughly 12,000 high-dimensional shape phenotypes from 10 multi-class datasets using the same algorithms and their ensembles. Fig. 2b displays the distribution of F1 scores and balanced accuracies for the top 10 base learners and select ensembles among multi-class datasets (see Table S3 and Fig. S5 for all algorithms). For base learners, the top 10 average F1 scores in descending order were obtained by mda, glm, penalized discriminant analysis (pda), linear discriminant analysis (lda), sparseLDA, *k*-nearest neighbors (kknn), nnet, multivariate adaptive regression splines (earth), stepwise linear discriminant analysis (stepLDA), and ranger (i.e., a rf variant). The top 10 and top five glm ensembles, as well as the top three rf ensemble, ranked first, second, and third with 82.0%, 81.2%, and 81.0% average F1 scores, respectively. In addition, the top three glm ensemble, alongside the top five and top 10 rf ensembles, tied for the fourth at 80.5% above mda, the leading base learner at 79.2%. Likewise, the top 10 base learners regarding balanced accuracy were mda, pda, lda, glm, sparseLDA, stepLDA, nnet, kknn, rf, and bagged multivariate adaptive regression splines (bagEarthGCV). The top 10 and top five glm ensembles tied for first with 88.4% average accuracies, whereas mda, the leading base learner, finished slightly behind at 88.3%. Just below this were the 88.0% to 88.2% accuracies achieved by the remaining ensembles.

To understand the composition and performance of the ensembles, we assessed the extent to which the base learner validation predictions deviated from the test predictions (Figs. 2c,d). For the top 10 base learners among binary datasets, we observed that the validation predictions exhibited 3.4%, 3.1%, 2.4%, 3.8%, 4.4%, and 6.0% decreases in F1, balanced accuracy, sensitivity, specificity, precision, and Kappa performance, respectively, compared to the test predictions (Fig. 2c). Conversely, the test and validation predictions for the top 10 base learners among multi-class datasets were nearly indistinguishable. While precision and Kappa were 1.5% and 1.1% higher, respectively, for the validation predictions, all other metrics showed mean differences of 0% to 0.1% (Fig. 2d).

Since model performance among datasets can be biased by poor or great performance within a minority of datasets, we also quantified relative model rank in terms of average F1 score and balanced accuracy within datasets (Fig. 3). Whereas a score of –1 indicates the lowest error or highest rank, 1 indicates the highest error or lowest rank. Much like the overall performance results, the top 10 average base learners within the binary datasets were rda, svmRadial, nnet, loclda, AdaBoost, hda, sparseLDA, mda, pda, and qda (Fig. 3a). The top three and top five rf ensembles finished first and second with –0.72 and –0.70 average ranks, respectively, above the – 0.61 of rda, the leading base learner. By contrast, the top 10 average base learners within the multi-class datasets diverged from the overall results. In descending order, they were loclda, qda, rda, mda, sparseLDA, pda, glm, lda, nnet, and hdda (Fig. 3b). The top ten and top five glm ensembles placed first and second with –0.80 and –0.73 average ranks, respectively, above the –0.58 of loclda, the leading base learner. Table S4 contains the full list of relative ranks.

**Figure 3.**
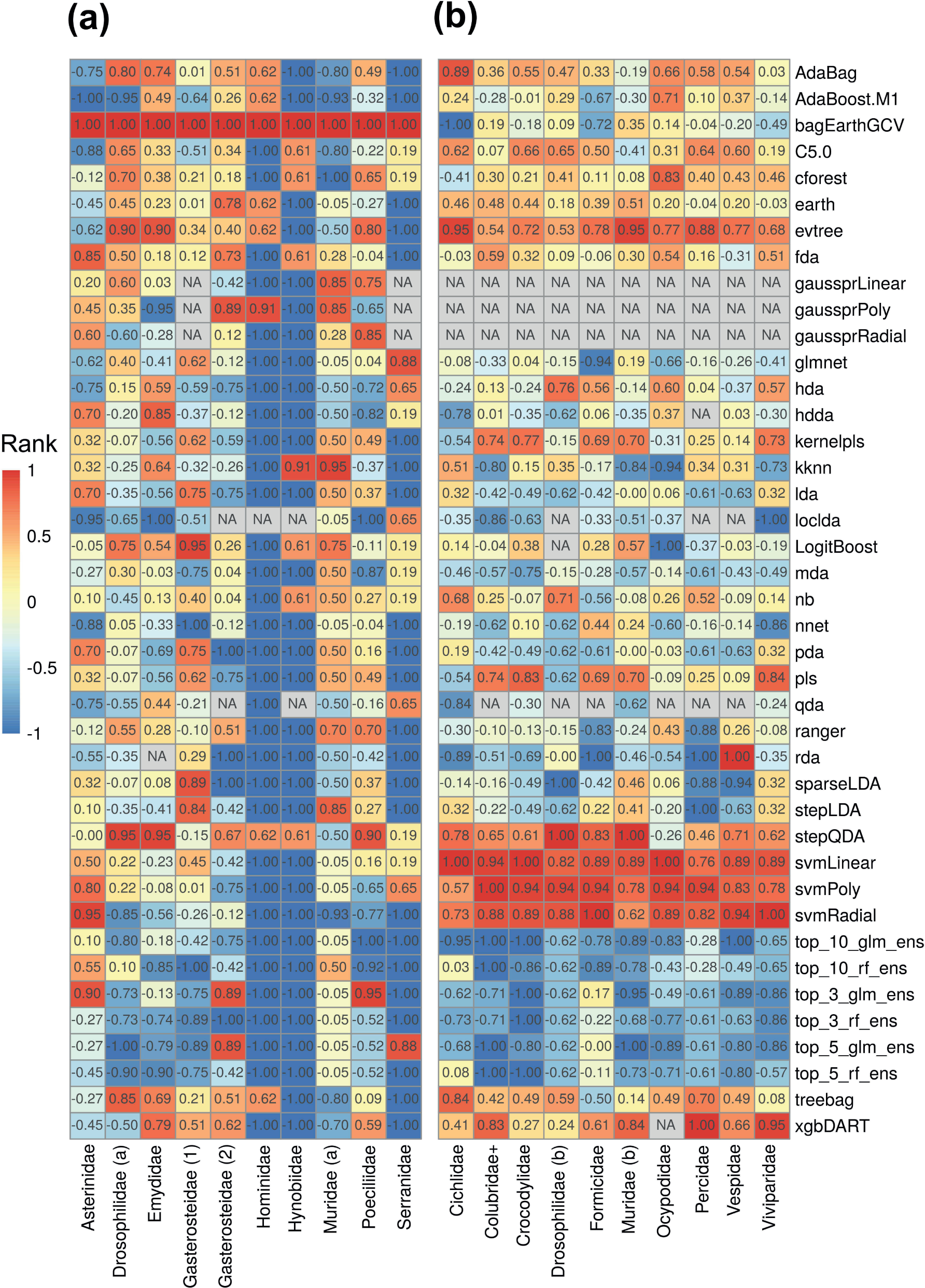
Relative method rank within (a) binary and (b) multi-class datasets according to average F1 and balanced accuracy score. Scores are normalized between –1 (lowest error, highest rank) and 1 (highest error, lowest rank). Datasets (columns) are listed alphabetically within classification task and methods are listed alphabetically overall.

To assess potential determinants of classification performance, we completed multiple regressions. Table 3 enumerates the means of the explanatory variables, alongside the F1 regression effect sizes for both the variable and task covariate. Here, effect refers to the average change in performance per unit increase in the variable: a unit for task is the change from binary to multi, whereas a unit for all continuous variables is 0.1, except for 0.01 in the case of shape distance and 1 for sample size.

**Table 3.** Summary of phenotypic and dataset variable means, effect sizes, and covariate task effect sizes in each F1 regression, as well as the model standard error (SE), F statistic, and overall R^2^.

| Variable | Binary | Multi | Variable Effect | Task Effect | SE | F | $R^2$ |
| --- | --- | --- | --- | --- | --- | --- | --- |
| $R^2$ | 0.2 | 0.5 | 5.5% | -36.2% | 13.6 | 9.0 | 0.46 |
| Shape distance | 0.06 | 0.08 | 2.1% | -23.8% | 14.1 | 7.8 | 0.42 |
| Covariance distance | 0.1 | 0.2 | -3.2% | -14.2% | 14.9 | 6.1 | 0.35 |
| Variance ratio | 1.2 | 1.5 | 0.4% | -20.9% | 15.7 | 4.7 | 0.28 |
| Class balance | 0.9 | 0.9 | -2.9% | -14.3% | 17.0 | 2.7 | 0.15 |
| Sample size | 981 | 1197 | 0% | -17.9% | 17.5 | 2.1 | 0.11 |

We found that R^2^ values derived from linear models of shape on class were the most predictive (Fig. 4a), followed by mean class shape distance (Fig. 4b), mean class covariance distance (Fig. 4c), and between– vs. within-class variance (Fig. 4d). Class balance (Fig. 4e) and total sample size (Fig. 4f) were substantially less predictive. Unsurprisingly, task was highly predictive in every model, resulting in 14.2% to 36.2% decreases in F1 as one moves from binary to multi-class classification. Balanced accuracy was influenced in the same manner, just to a lesser extent (Fig. S6). Table S5 describes the effect sizes and model fit measures for balanced accuracy.

**Figure 4.**
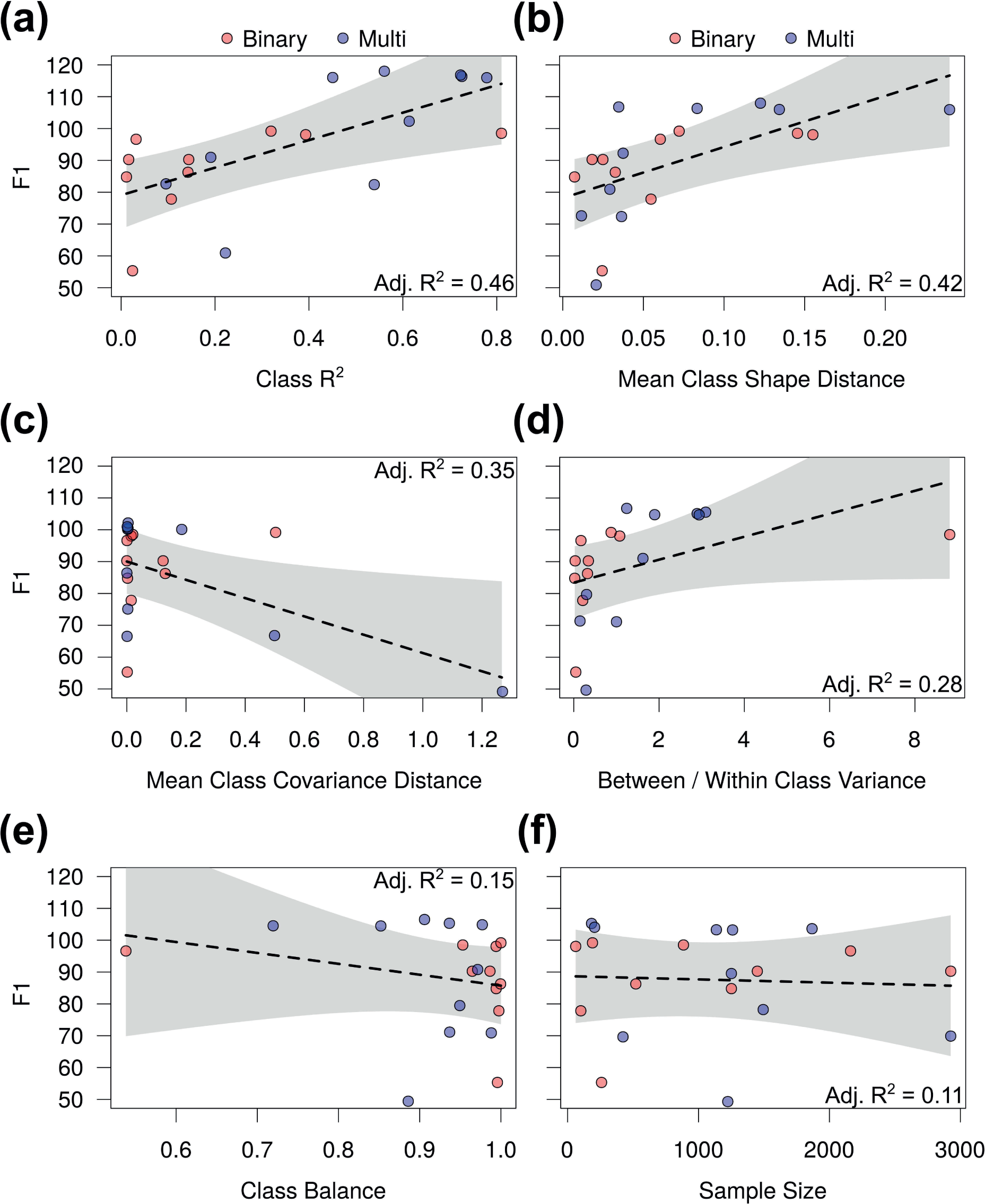
F1 multiple regression plots with classification task (red, binary; blue, multi) plus (a) class R2, (b) mean class shape distance, (c) mean class covariance distance, (d) between– vs. within-class variance, (e) class balance, or (f) sample size explanatory variables. Lines of best fit with 95% confidence intervals are shown alongside model R2 values.

## 4.0 Discussion

We have presented a large-scale empirical analysis of classification algorithms, alongside a generic ensemble learning framework for classifying high-dimensional phenotypes. Classification is a fundamental problem in biology that has seen renewed interest over the past five years due to the explosion of data and machine learning techniques. Unfortunately, most emphasis has been placed on developing methods for a particular classification task or on optimizing and comparing a small set of learning algorithms for a specific phenotypic dataset. Our first aim was to quantify average method performance across high-dimensional shape datasets with different anatomies, variance-covariance patterns, mean distances, class distributions, and sample sizes. Our second aim was to combine these learners into a stronger phenotypic learner via ensemble learning and compare its performance. This culminated in the *pheble* R package, which offers a flexible, effective, and streamlined solution for classifying high-dimensional phenotypes. The workflow contains functions for preprocessing, training and strategically stacking a multitude of models to build an ensemble, and model evaluation.

To preprocess each dataset, we implemented an 85/15% training/test split with 15% of the training data reserved for validation. This split ratio appeared sufficient on average, given the final ensemble results; however, these percentages are merely a guideline as they are a function of the minimum class sample size. For example, while the multi-class validation and test predictions were nearly equivalent, the binary validation predictions were notably worse than the test predictions, suggesting that more data were needed. We discovered that the *Hominidae* sacrum dataset was single-handedly driving this difference. While all other datasets exhibited average validation and test performance deviations between 0% and 8%, the *Hominidae* dataset displayed average deviations of 25% (Table S6). Being the second smallest dataset with *N*=101 observations, the holdout validation set was limited to *N*=13 observations, so any classification errors in these data would be magnified. Larger validation and test set partitions are therefore recommended in similar scenarios. Interestingly, even with errant validation predictions, the *Hominidae* ensembles managed to achieve test performance on par with all other methods, likely because its class boundaries were easily separable. Small datasets with less distinct classes may not be so fortunate.

The other core preprocessing steps were anomaly detection and dimensionality reduction. Since this study dealt with Procrustes shape coordinates, we opted for the classical Procrustes distance solution to remove outliers. Nevertheless, we also introduced a more generic approach to make the classification pipeline end-to-end. We showed that anomaly scores from autoencoders and extended isolation forests are highly correlated with Procrustes distance to the mean, with the former being the superior option. Importantly, though, these methods learn specific features of a dataset as opposed to an aggregate feature metric, like distance, so the comparison is imperfect. Merging domain-specific and learning-based anomaly scores may offer the most insight, but more exploration is needed. In terms of dimensionality reduction, we provide PCA and autoencoder options for linear and non-linear decompositions, respectively, although other extracted features or even the raw data can be used. We ultimately chose PCs due to the highly correlated and Euclidean nature of Procrustes coordinates projected into tangent space.

Existing R packages for ensemble learning either have a limited pool of classification algorithms or are unable to train multi-class ensembles. Since ensemble models are most effective when they incorporate many diverse base learners (LeDell, 2015; van der Laan et al., 2007), we exploited the enormously successful and comprehensive training interface of *caret*. After screening each algorithm for errors and overall feasibility (e.g., performance and training time), we selected 33 learners for binary classification and 30 learners for multi-class classification. We experimented with ensembles that stacked the top three, top five, and top 10 base learners, but we suspect that more learners could improve performance. The top base learners and ensembles among datasets but within classification task were fairly consistent between our overall performance metrics, F1 and balanced accuracy. In descending order, the best binary class approaches were the top three rf ensemble, rda, top five rf ensemble, top 10 glm ensemble, hda, nnet, loclda, top 10 rf ensemble, mda, svmPoly, AdaBoost, svmRadial, sparseLDA, and pls. Likewise, the best multi-class methods were the top three, 10, and five glm ensembles, top five and 10 rf ensembles, mda, top three rf ensemble, pda, glmnet, lda, sparseLDA, nnet, stepLDA, kknn, bagEarthGCV, and ranger. Relative to the top ranked base learner, the best binary class ensemble improved average F1 performance by 0.4%, while the best multi-class ensemble improved average F1 performance by 3%.

We additionally evaluated algorithm performance within datasets to avoid particular class biases. Some methods, for example, may be highly effective or ineffective for a specific dataset and this could therefore inflate or deflate their overall performance. We found that the top base learners varied from dataset to dataset, whereas the ensembles consistently achieved superior performance. Such variability is not surprising, given that phenotypic spaces and class boundaries vary among datasets. But this result is critical to underscore, because it clearly shows that one cannot rely on the performance results from other studies to inform a new, unrelated study. Deploying an ensemble, on the other hand, will ensure effective, reliable classification. Even if the ensemble does not finish atop the base learners, the user can easily discover the best model and retrieve it thanks to the ensemble process. Another point worth mentioning is we only evaluated glm and rf metalearners due to their robustness to overfitting. The rf metalearners greatly outperformed glms for binary classification and vice versa, albeit to a much lesser extent, for multi-class classification. Considering the range of the binary classification metalearner results, we recommend experimenting with alternatives, especially since that functionality is supported.

Our final aim was to quantify the impact of various dataset and phenotypic properties on classification performance. Using classification task as a covariate, we found that in each multiple regression model, task explained the highest proportion of variance in F1 and balanced accuracy. This was expected and merely indicates that multi-class performance is lower on average than binary performance. We additionally observed that higher class R^2^, mean class shape distance, and between– vs. within-class variance values increased performance. Computing each of these measures essentially involves maximizing differences among classes, so the positive associations make sense. By contrast, increases in covariance distance decreased performance. Because this measure reflects differences in the shape of the covariance matrix between classes, we can conclude that learning algorithms struggle with increasingly disparate class distributions on average. Balance in the number of observations per class appeared to decrease performance, but this model exhibited high error and surely reflects noise, as class imbalances are a known problem for many learning-based models (Sun et al., 2009). The equally poor predictive power of sample size suggests that our base learners and ensembles can support smaller samples. However, the smallest samples were easily discriminated by most methods, so this result should be interpreted with caution.

## 5.0 Conclusions

Learning-based classification is a complex task driven by many hyperparameters. We introduced the R package *pheble* to perform a meta-analysis of classification algorithms and provide a streamlined ensemble learning workflow for classifying high-dimensional phenotypes. Binary and multi-class classification tasks relevant to evolutionary biology, developmental biology, and ecology were considered. In total, we classified over 20,000 high-dimensional shape phenotypes using 33 algorithms and their ensembles. We found that discriminant analysis variants and neural networks were the most accurate learners on average. However, there was considerable variability in base learner performance between datasets. Ensemble models, on the other hand, achieved the highest performance on average, both within and among datasets. By quantifying the extent to which certain dataset and phenotypic properties influence these models, we also offer likely explanations for variation in performance. Researchers interested in maximizing classification performance stand to benefit from the simplicity and effectiveness of our approach.

## Acknowledgements

We would like to acknowledge the generous funding from a CIHR Foundation Grant (#159920), an NSERC Discovery Grant (#238992-17), and an NIH R01 Grant (#2R01DE019638).

## Author Contributions

J.D. and B.H. conceived the ideas. J.D. performed the meta-analysis, wrote the R package, and wrote the first draft. All authors discussed aspects of the research and contributed to writing and revising the paper.

## Data Availability

We used 20 publicly available datasets and referred to them by family: *Asterinidae* (Araújo et al., 2014), *Drosophilidae* (a/b) (Pitchers et al., 2013), *Emydidae* (Stayton et al., 2018), *Gasterosteidae* (1) (Schutz et al., 2022), *Gasterosteidae* (2) (Fraser & El-Sabaawi, 2022), *Hominidae* (Krenn et al., 2022), *Hynobiidae* (Jia et al., 2022), *Muridae* (a/b) (Devine et al., 2022), *Poeciliidae* (Riesch et al., 2016), Serranidae (Alós et al., 2014), *Cichlidae* (Ronco et al., 2020), *Colubridae*+ (Head & Polly, 2015), *Crocodylidae* (Watanabe & Slice, 2014), *Formicidae* (Kennedy et al., 2014), *Ocypodidae* (Hopkins et al., 2016), *Percidae* (Martin & Mendelson, 2014), *Vespidae* (Perrard et al., 2014), and *Viviparidae* (Van Bocxlaer & Hunt, 2013). Dataset details are listed in Table S1. The data and code to reproduce our analysis are available at doi.org/10.5281/zenodo.7949383. The R package code is available at github.com/jaydevine/pheble.

