## Supplementary Info for "Classifying high-dimensional phenotypes with ensemble learning"

***Table S1.*** Complete summary of phenotypic datasets, including the family, landmarked anatomy, total sample size (*N*), class, number of class levels, number of landmarks (*p*), the dimensionality (*k*), and the original data repository.

| Family | Anatomy | *N* | Class | Levels | *p* | *k* | Data Repository |
|---|---|---|---|---|---|---|---|
| Asterinidae | Body | 885 | Sex | 2 | 10 | 2 | http://dx.doi.org/10.5061/dryad.4h31p |
| Drosophilidae (a) | Wing | 2926 | Sex | 2 | 48 | 2 | http://dx.doi.org/10.5061/dryad.r43k1.2 |
| Emydidae | Shell | 2161 | Habitat | 2 | 53 | 3 | https://doi.org/10.5061/dryad.8r0m76h |
| Gasterosteidae (1) | Skull | 190 | Habitat | 2 | 70 | 3 | https://doi.org/10.5061/dryad.xd2547dkw |
| Gasterosteidae (2) | Body | 521 | Sex | 2 | 15 | 2 | https://doi.org/10.5061/dryad.bzkh189cx |
| Hominidae | Sacrum | 101 | Sex | 2 | 100 | 3 | https://doi.org/10.1371/journal.pone.0264770 |
| Hynobiidae/Cryptobranchidae | Palate | 62 | Habitat | 2 | 24 | 2 | https://doi.org/10.5061/dryad.c59zw3r8x |
| Muridae (a) | Cranium | 1251 | Sex | 2 | 844 | 3 | https://doi.org/10.1038/s41597-022-01338-x |
| Poeciliidae | Body | 1449 | Sex | 2 | 13 | 2 | https://doi.org/10.5061/dryad.2sh53 |
| Serranidae/Sparidae | Body | 259 | Site | 2 | 13 | 2 | https://doi.org/10.5061/dryad.1k571 |
| Cichlidae | Jaw | 1136 | Tribe | 14 | 42 | 3 | https://doi.org/10.5061/dryad.9w0vt4bbf |
| Colubridae+ | Vertebrae | 1260 | Species | 15 | 12 | 2 | https://doi.org/10.5061/dryad.jq285 |
| Crocodylidae/Alligatoridae | Cranium | 183 | Species | 8 | 78 | 3 | http://dx.doi.org/10.5061/dryad.14fn1 |
| Drosophilidae (b) | Wing | 2926 | Elevation | 9 | 48 | 2 | http://dx.doi.org/10.5061/dryad.r43k1.2 |
| Formicidae | Face | 1494 | Species | 6 | 11 | 2 | http://dx.doi.org/10.5061/dryad.f65bn |
| Muridae (b) | Cranium | 1251 | Genotype | 26 | 844 | 3 | https://doi.org/10.1038/s41597-022-01338-x |
| Ocypodidae | Carapace | 1867 | Species | 16 | 21 | 2 | http://dx.doi.org/10.5061/dryad.vv197 |
| Percidae | Body | 423 | Species | 15 | 10 | 2 | https://doi.org/10.5061/dryad.n28rf |
| Vespidae | Wing | 206 | Species | 8 | 19 | 2 | http://dx.doi.org/10.5061/dryad.4588r |
| Viviparidae | Shell | 1224 | Population | 22 | 127 | 2 | https://doi.org/10.5061/dryad.vm523 |

***Table S2.*** Summary of performance metrics for each method among all binary class datasets.

| Method | F1 | Accuracy | Sensitivity | Specificity | Precision | Kappa |
|---|---|---|---|---|---|---|
| AdaBag | 0.86 | 0.84 | 0.88 | 0.81 | 0.85 | 0.7 |
| AdaBoost.M1 | 0.9 | 0.88 | 0.9 | 0.87 | 0.9 | 0.78 |
| bagEarthGCV | NA | 0.13 | NA | NA | 0.14 | NA |
| C5.0 | 0.88 | 0.87 | 0.89 | 0.85 | 0.88 | 0.75 |
| cforest | 0.88 | 0.86 | 0.9 | 0.81 | 0.87 | 0.73 |
| earth | 0.85 | 0.85 | 0.85 | 0.84 | 0.85 | 0.69 |
| evtree | 0.84 | 0.81 | 0.87 | 0.75 | 0.83 | 0.63 |
| fda | 0.84 | 0.84 | 0.83 | 0.85 | 0.86 | 0.69 |
| gaussprLinear | 0.85 | 0.86 | 0.83 | 0.89 | 0.88 | 0.68 |
| gaussprPoly | 0.79 | 0.77 | 0.74 | 0.8 | 0.84 | 0.55 |
| gaussprRadial | 0.86 | 0.83 | 0.83 | 0.83 | 0.9 | 0.68 |
| glmnet | 0.87 | 0.83 | 0.89 | 0.77 | 0.86 | 0.67 |
| hda | 0.9 | 0.9 | 0.89 | 0.91 | 0.92 | 0.8 |
| hdda | 0.87 | 0.87 | 0.87 | 0.86 | 0.87 | 0.72 |
| kernelpls | 0.89 | 0.87 | 0.9 | 0.85 | 0.88 | 0.75 |
| kknn | 0.86 | 0.85 | 0.89 | 0.82 | 0.86 | 0.71 |
| lda | 0.89 | 0.88 | 0.88 | 0.87 | 0.9 | 0.76 |
| loclda | 0.9 | 0.9 | 0.89 | 0.9 | 0.91 | 0.79 |
| LogitBoost | 0.86 | 0.84 | 0.86 | 0.82 | 0.86 | 0.68 |
| mda | 0.89 | 0.89 | 0.88 | 0.9 | 0.9 | 0.78 |
| nb | 0.88 | 0.87 | 0.87 | 0.86 | 0.88 | 0.74 |
| nnet | 0.9 | 0.89 | 0.91 | 0.88 | 0.9 | 0.79 |
| pda | 0.89 | 0.88 | 0.89 | 0.86 | 0.89 | 0.76 |
| pls | 0.89 | 0.88 | 0.89 | 0.87 | 0.9 | 0.77 |
| qda | 0.89 | 0.88 | 0.88 | 0.87 | 0.9 | 0.76 |
| ranger | 0.87 | 0.84 | 0.88 | 0.8 | 0.86 | 0.7 |
| rda | 0.91 | 0.91 | 0.91 | 0.91 | 0.91 | 0.8 |
| sparseLDA | 0.89 | 0.88 | 0.9 | 0.86 | 0.89 | 0.76 |
| stepLDA | 0.89 | 0.87 | 0.88 | 0.87 | 0.89 | 0.76 |
| stepQDA | 0.82 | 0.78 | 0.83 | 0.73 | 0.82 | 0.57 |
| svmLinear | 0.89 | 0.88 | 0.88 | 0.87 | 0.89 | 0.76 |
| svmPoly | 0.89 | 0.9 | 0.87 | 0.92 | 0.92 | 0.8 |
| svmRadial | 0.89 | 0.89 | 0.89 | 0.88 | 0.89 | 0.78 |
| top_10_glm_ens | 0.9 | 0.9 | 0.9 | 0.91 | 0.92 | 0.81 |
| top_10_rf_ens | 0.9 | 0.89 | 0.89 | 0.89 | 0.9 | 0.79 |
| top_3_glm_ens | 0.85 | 0.81 | 0.76 | 0.85 | 0.94 | 0.62 |
| top_3_rf_ens | 0.91 | 0.91 | 0.92 | 0.9 | 0.91 | 0.82 |
| top_5_glm_ens | 0.85 | 0.81 | 0.82 | 0.81 | 0.87 | 0.63 |
| top_5_rf_ens | 0.91 | 0.9 | 0.9 | 0.91 | 0.91 | 0.81 |
| treebag | 0.86 | 0.84 | 0.88 | 0.8 | 0.85 | 0.69 |

| | | | | | | |
|---|---|---|---|---|---|---|
| xgbDART | 0.86 | 0.85 | 0.86 | 0.84 | 0.88 | 0.7 |

***Table S3.*** Summary of performance metrics for each method among all multi-class datasets.

| **Method** | **F1** | **Accuracy** | **Sensitivity** | **Specificity** | **Precision** | **Kappa** |
|---|---|---|---|---|---|---|
| AdaBag | 0.68 | 0.8 | 0.64 | 0.97 | 0.72 | 0.67 |
| AdaBoost.M1 | 0.73 | 0.84 | 0.69 | 0.98 | 0.74 | 0.72 |
| bagEarthGCV | 0.76 | 0.85 | 0.72 | 0.98 | 0.77 | 0.73 |
| C5.0 | 0.68 | 0.81 | 0.65 | 0.97 | 0.7 | 0.67 |
| cforest | 0.72 | 0.81 | 0.64 | 0.97 | 0.73 | 0.68 |
| earth | 0.71 | 0.82 | 0.67 | 0.97 | 0.73 | 0.69 |
| evtree | 0.6 | 0.74 | 0.52 | 0.96 | 0.58 | 0.55 |
| fda | 0.72 | 0.83 | 0.69 | 0.97 | 0.73 | 0.7 |
| glmnet | 0.76 | 0.86 | 0.74 | 0.98 | 0.77 | 0.75 |
| hda | 0.71 | 0.83 | 0.69 | 0.97 | 0.73 | 0.7 |
| hdda | 0.73 | 0.84 | 0.71 | 0.98 | 0.73 | 0.71 |
| kernelpls | 0.68 | 0.78 | 0.59 | 0.97 | 0.73 | 0.66 |
| kknn | 0.76 | 0.85 | 0.73 | 0.98 | 0.76 | 0.73 |
| lda | 0.76 | 0.86 | 0.74 | 0.98 | 0.76 | 0.75 |
| loclda | 0.73 | 0.84 | 0.71 | 0.98 | 0.72 | 0.71 |
| LogitBoost | 0.73 | 0.82 | 0.67 | 0.97 | 0.72 | 0.69 |
| mda | 0.79 | 0.88 | 0.78 | 0.98 | 0.79 | 0.78 |
| nb | 0.7 | 0.83 | 0.68 | 0.97 | 0.73 | 0.69 |
| nnet | 0.76 | 0.85 | 0.73 | 0.98 | 0.76 | 0.73 |
| pda | 0.76 | 0.86 | 0.74 | 0.98 | 0.76 | 0.75 |
| pls | 0.68 | 0.78 | 0.59 | 0.97 | 0.74 | 0.66 |
| ranger | 0.75 | 0.85 | 0.72 | 0.98 | 0.79 | 0.74 |
| rda | 0.71 | 0.84 | 0.7 | 0.98 | 0.71 | 0.69 |
| sparseLDA | 0.76 | 0.86 | 0.74 | 0.98 | 0.76 | 0.75 |
| stepLDA | 0.75 | 0.86 | 0.74 | 0.98 | 0.75 | 0.74 |
| stepQDA | 0.58 | 0.74 | 0.52 | 0.96 | 0.58 | 0.51 |
| svmLinear | 0.51 | 0.68 | 0.41 | 0.94 | 0.45 | 0.34 |
| svmPoly | 0.52 | 0.68 | 0.41 | 0.95 | 0.45 | 0.35 |
| svmRadial | 0.54 | 0.68 | 0.42 | 0.95 | 0.46 | 0.37 |
| top_10_glm_ens | 0.82 | 0.88 | 0.79 | 0.98 | 0.8 | 0.79 |
| top_10_rf_ens | 0.8 | 0.88 | 0.78 | 0.98 | 0.8 | 0.78 |
| top_3_glm_ens | 0.81 | 0.88 | 0.78 | 0.98 | 0.81 | 0.78 |
| top_3_rf_ens | 0.81 | 0.88 | 0.78 | 0.98 | 0.79 | 0.78 |
| top_5_glm_ens | 0.81 | 0.88 | 0.79 | 0.98 | 0.83 | 0.79 |
| top_5_rf_ens | 0.8 | 0.88 | 0.78 | 0.98 | 0.79 | 0.79 |
| treebag | 0.68 | 0.8 | 0.62 | 0.97 | 0.68 | 0.65 |
| xgbDART | 0.65 | 0.74 | 0.52 | 0.96 | 0.62 | 0.56 |

***Table S4.*** Average relative rank within datasets.

| Method | Binary | Multi |
|---|---|---|
| AdaBag | -0.04 | 0.42 |
| AdaBoost.M1 | -0.45 | 0.03 |
| bagEarthGCV | 1 | -0.19 |
| C5.0 | -0.13 | 0.38 |
| cforest | 0.08 | 0.28 |
| earth | -0.07 | 0.28 |
| evtree | 0.08 | 0.76 |
| fda | 0.12 | 0.21 |
| gaussprLinear | 0 | NA |
| gaussprPoly | 0.11 | NA |
| gaussprRadial | -0.13 | NA |
| glmnet | -0.13 | -0.28 |
| hda | -0.39 | 0.17 |
| hdda | -0.23 | -0.21 |
| kernelpls | -0.23 | 0.3 |
| kknn | -0.04 | -0.18 |
| lda | -0.23 | -0.25 |
| loclda | -0.5 | -0.58 |
| LogitBoost | 0.29 | -0.03 |
| mda | -0.29 | -0.44 |
| nb | 0.08 | 0.17 |
| nnet | -0.54 | -0.24 |
| pda | -0.26 | -0.29 |
| pls | -0.25 | 0.29 |
| qda | -0.26 | -0.5 |
| rf | -0.05 | -0.2 |
| rda | -0.61 | -0.44 |
| sparseLDA | -0.29 | -0.32 |
| stepLDA | -0.21 | -0.19 |
| stepQDA | 0.42 | 0.64 |
| svmLinear | -0.12 | 0.91 |
| svmPoly | -0.18 | 0.87 |
| svmRadial | -0.55 | 0.87 |
| top_10_glm_ens | -0.61 | -0.8 |
| top_10_rf_ens | -0.5 | -0.6 |
| top_3_glm_ens | -0.19 | -0.66 |
| top_3_rf_ens | -0.72 | -0.68 |
| top_5_glm_ens | -0.38 | -0.73 |
| top_5_rf_ens | -0.7 | -0.61 |
| treebag | -0.01 | 0.37 |
| xgbDART | -0.21 | 0.64 |

***Table S5.*** Summary of phenotypic and dataset variable means, effect sizes, and covariate task effect sizes in each balanced accuracy regression, as well as the model standard error (SE), F statistic, and overall $R^2$.

| **Variable** | **Binary** | **Multi** | **Variable Effect** | **Task Effect** | **SE** | **F** | **$R^2$** |
|---|---|---|---|---|---|---|---|
| $R^2$ | 0.2 | 0.5 | 4.1% | -17.2% | 9.1 | 7.7 | 0.41 |
| Shape distance | 0.06 | 0.08 | 1.4% | -7.4% | 10.1 | 4.8 | 0.29 |
| Covariance distance | 0.1 | 0.2 | -2.1% | NA | 10.8 | 4.9 | 0.17 |
| Variance ratio | 1.2 | 1.5 | 0.3% | -6.2% | 10.8 | 3.0 | 0.17 |
| Class balance | 0.9 | 0.9 | -0.5% | NA | 12.2 | 0 | -0.05 |
| Sample size | 981 | 1197 | 0% | NA | 12.2 | 0.1 | -0.05 |

***Table S6.*** Summary of average validation and test performance among all methods but within datasets.

| **Dataset** | **Task** | **Vali F1** | **Test F1** | **Vali Accuracy** | **Test Accuracy** |
|---|---|---|---|---|---|
| Asterinidae | Binary | 0.99 | 0.99 | 0.95 | 0.97 |
| Drosophilidae (a) | Binary | 0.85 | 0.90 | 0.87 | 0.91 |
| Emydidae | Binary | 0.94 | 0.97 | 0.82 | 0.85 |
| Gasterosteidae (1) | Binary | 1.00 | 0.99 | 0.97 | 0.99 |
| Gasterosteidae (2) | Binary | 0.90 | 0.86 | 0.91 | 0.87 |
| Hominidae | Binary | 0.53 | 0.78 | 0.54 | 0.79 |
| Hynobiidae | Binary | 0.96 | 0.98 | 0.91 | 0.97 |
| Muridae (a) | Binary | 0.84 | 0.85 | 0.82 | 0.82 |
| Poeciliidae | Binary | 0.89 | 0.90 | 0.85 | 0.86 |
| Serranidae | Binary | 0.49 | 0.55 | 0.55 | 0.60 |
| Cichlidae | Multi | 0.82 | 0.87 | 0.88 | 0.92 |
| Colubridae+ | Multi | 0.84 | 0.87 | 0.89 | 0.92 |
| Crocodylidae | Multi | 0.86 | 0.89 | 0.89 | 0.91 |
| Drosophilidae (b) | Multi | 0.49 | 0.54 | 0.70 | 0.73 |
| Formicidae | Multi | 0.63 | 0.62 | 0.78 | 0.77 |
| Muridae (b) | Multi | 0.73 | 0.74 | 0.84 | 0.85 |
| Ocypodidae | Multi | 0.88 | 0.88 | 0.93 | 0.93 |
| Percidae | Multi | 0.62 | 0.54 | 0.76 | 0.72 |
| Vespidae | Multi | 0.82 | 0.88 | 0.89 | 0.92 |
| Viviparidae | Multi | 0.34 | 0.32 | 0.55 | 0.56 |

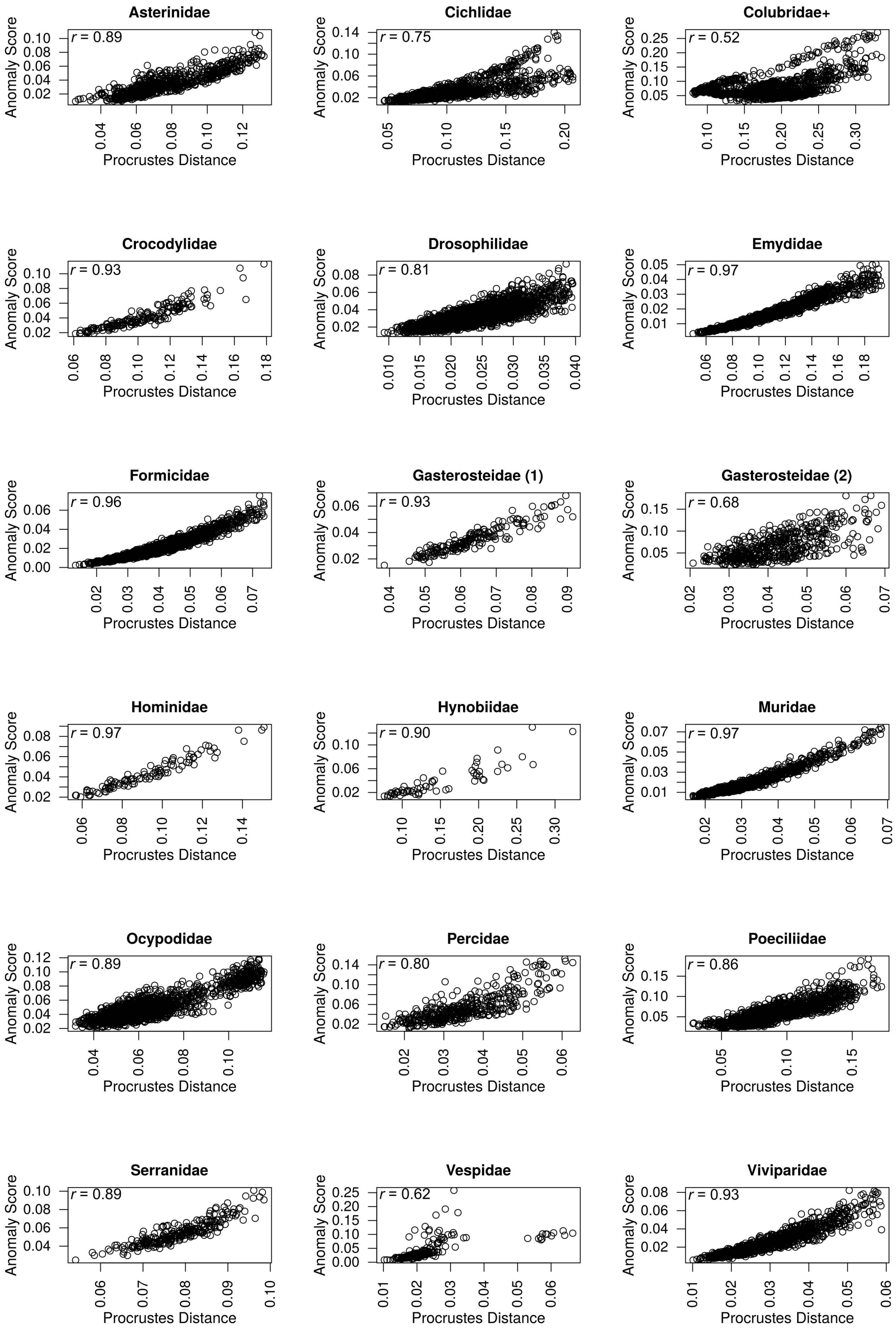


**Figure S1**. Bivariate plots for autoencoder anomaly scores and Procrustes distances to the mean. Correlation coefficients (*r*) are included. Plots are organized alphabetically by dataset.

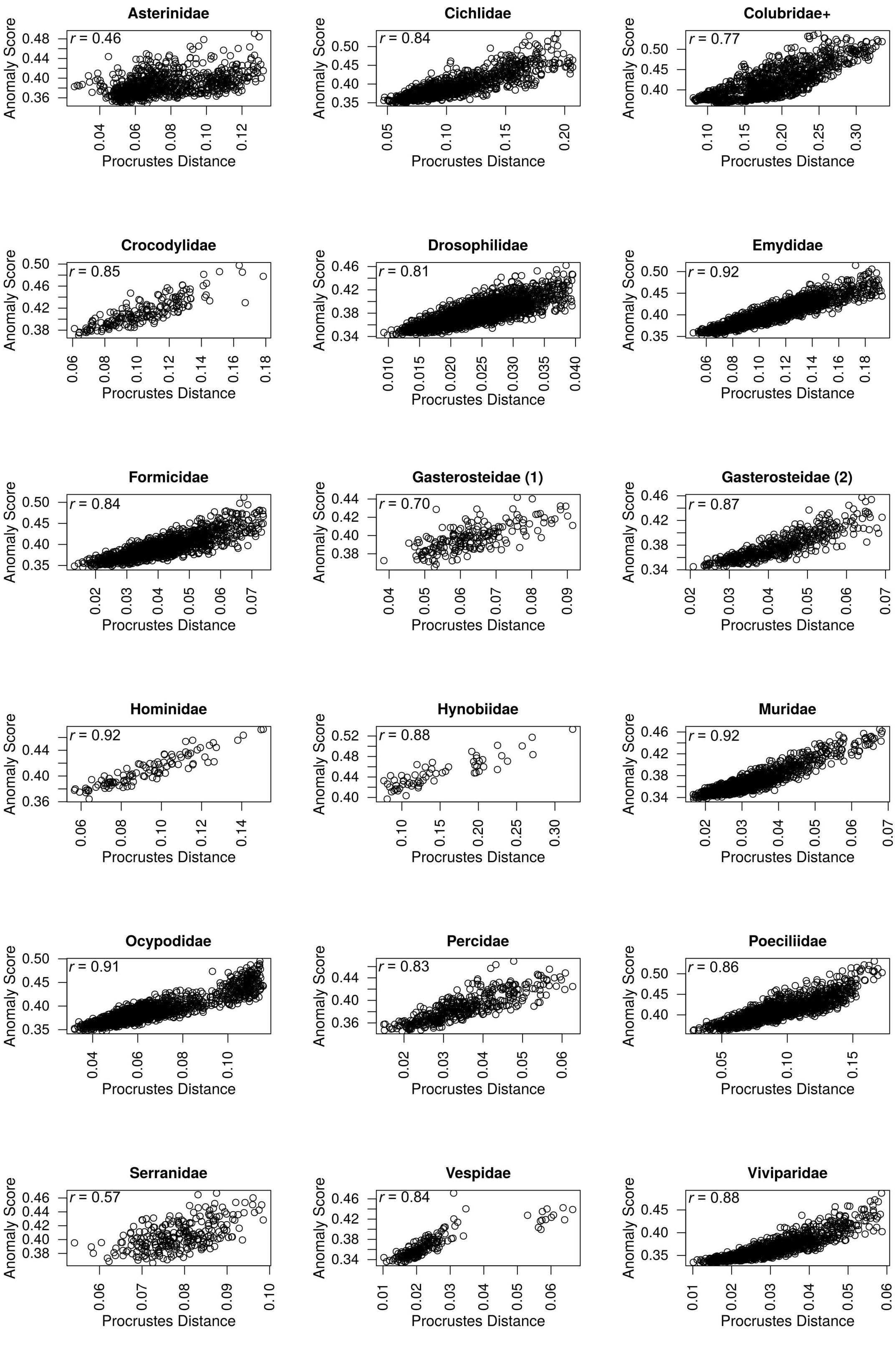


**Figure S2**. Bivariate plots for extended isolation forest anomaly scores and Procrustes distances to the mean. Correlation coefficients (*r*) are included. Plots are organized alphabetically by dataset.

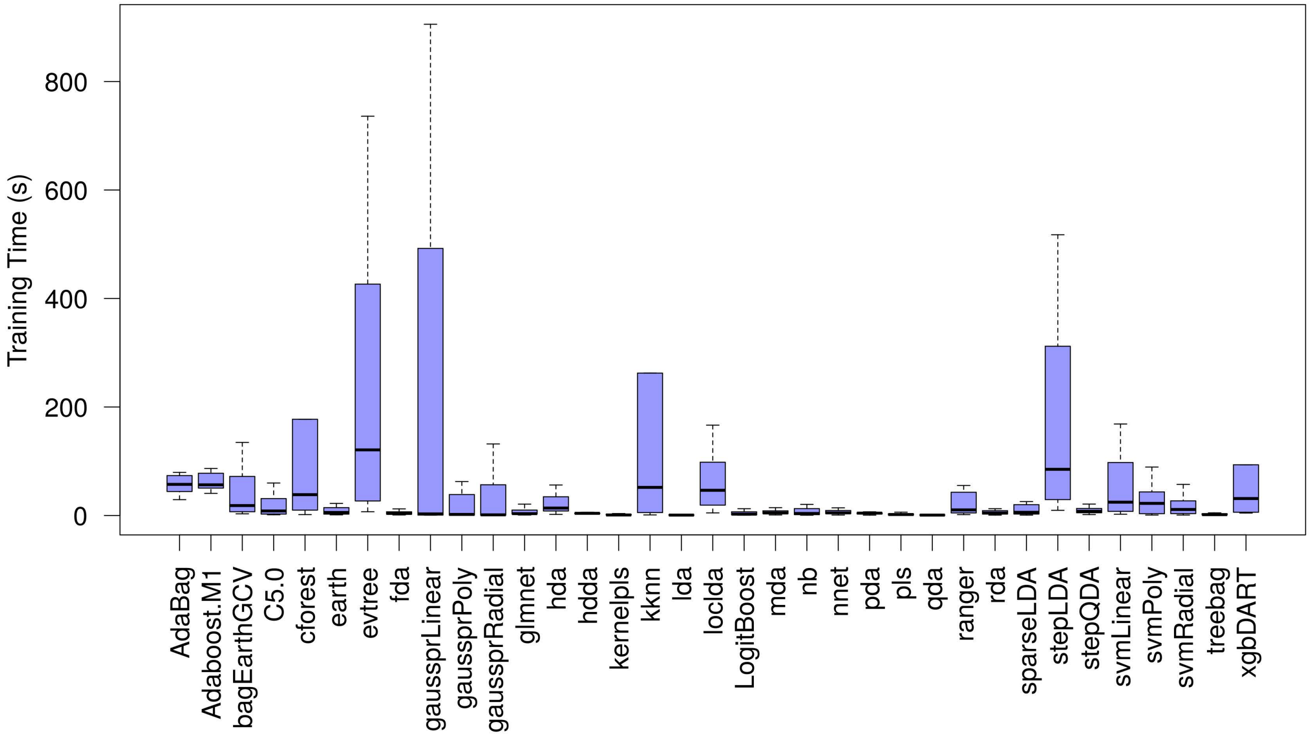


**Figure S3**. Distribution of training times (in seconds) for each method ordered alphabetically.

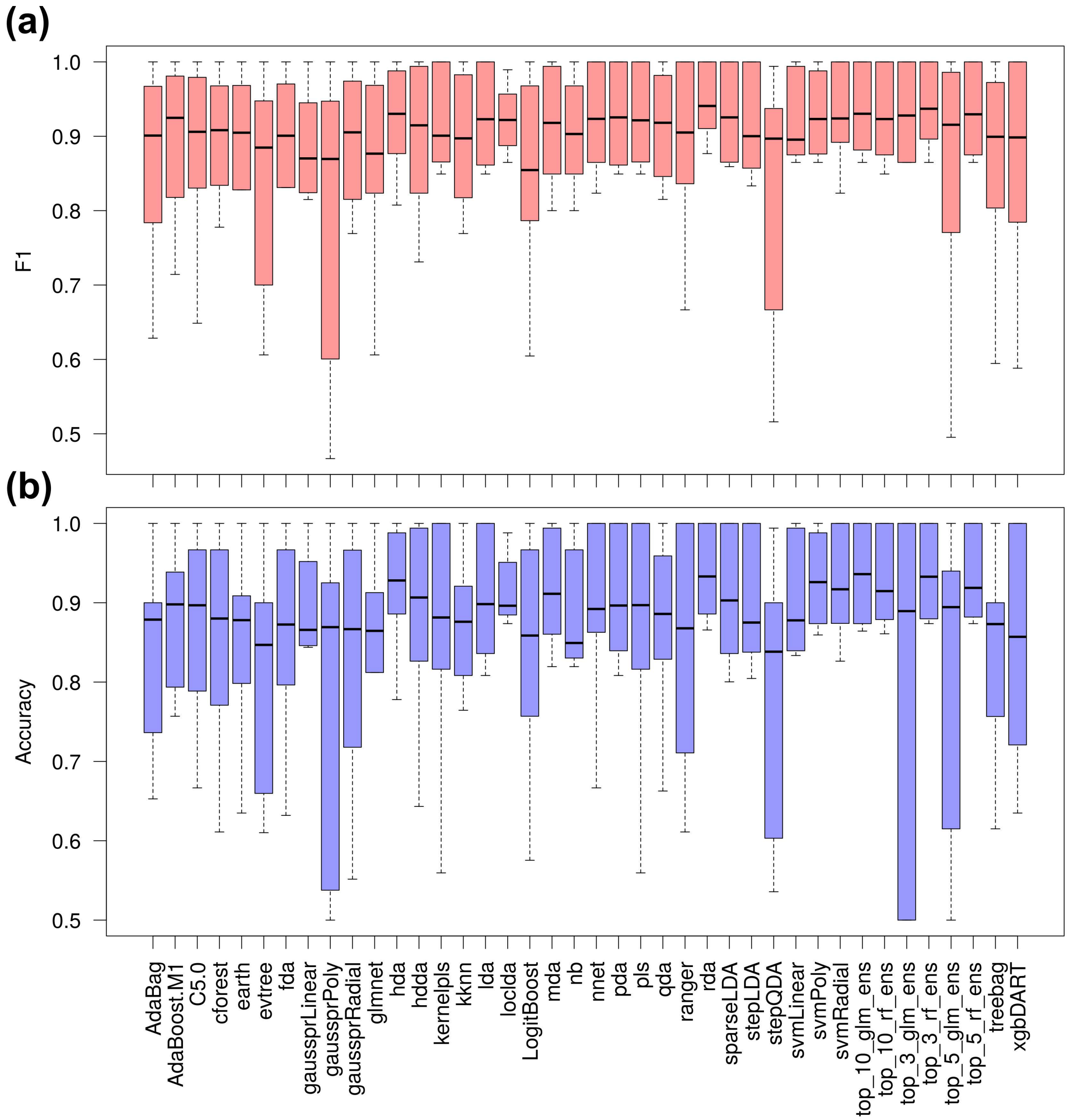


**Figure S4**. Distribution of (a) F1 and (b) balanced accuracy scores for every method among binary class datasets. Methods are ordered alphabetically.

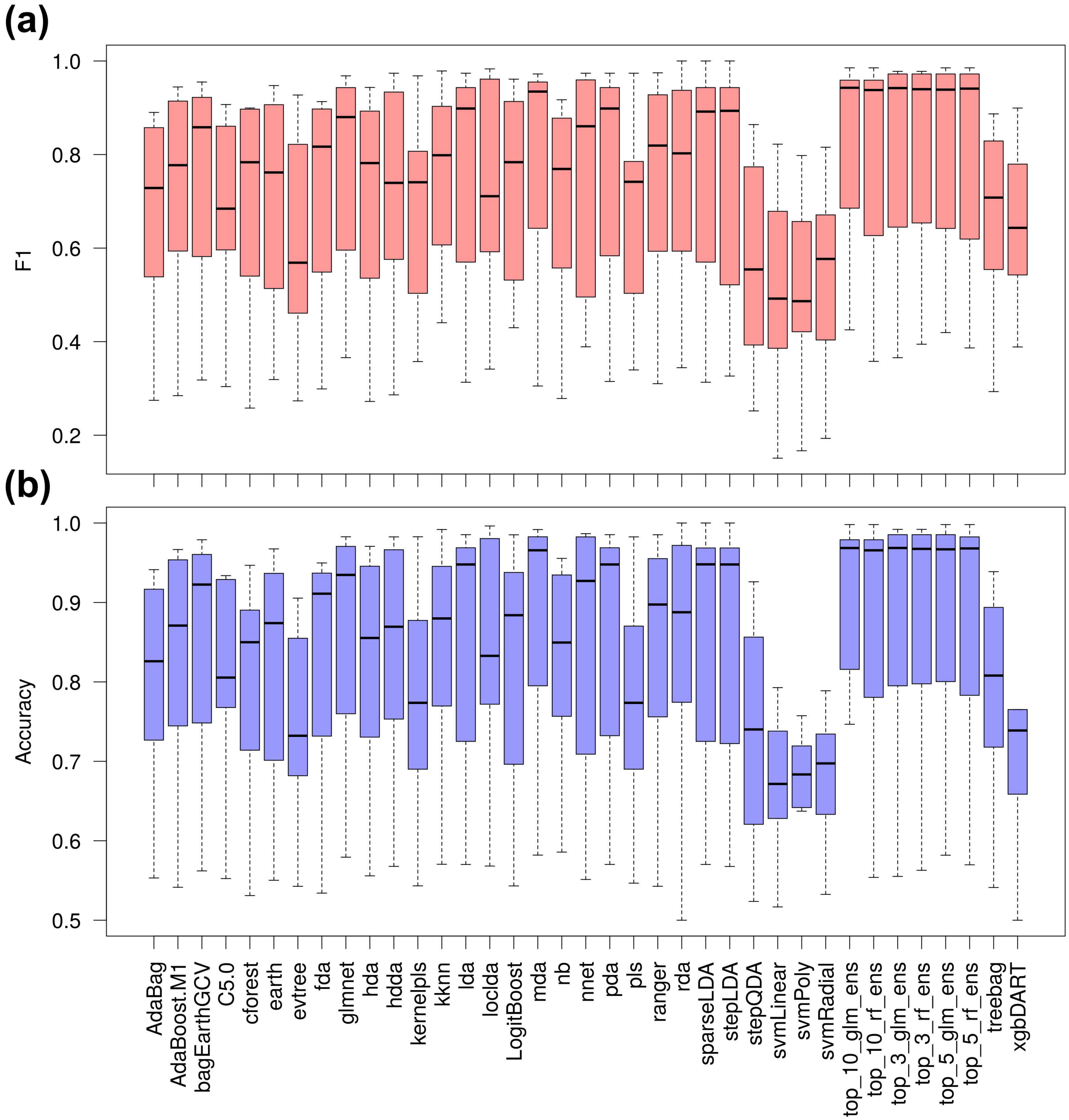


**Figure S5**. Distribution of (a) F1 and (b) balanced accuracy scores for every method among multi-class datasets. Methods are ordered alphabetically.

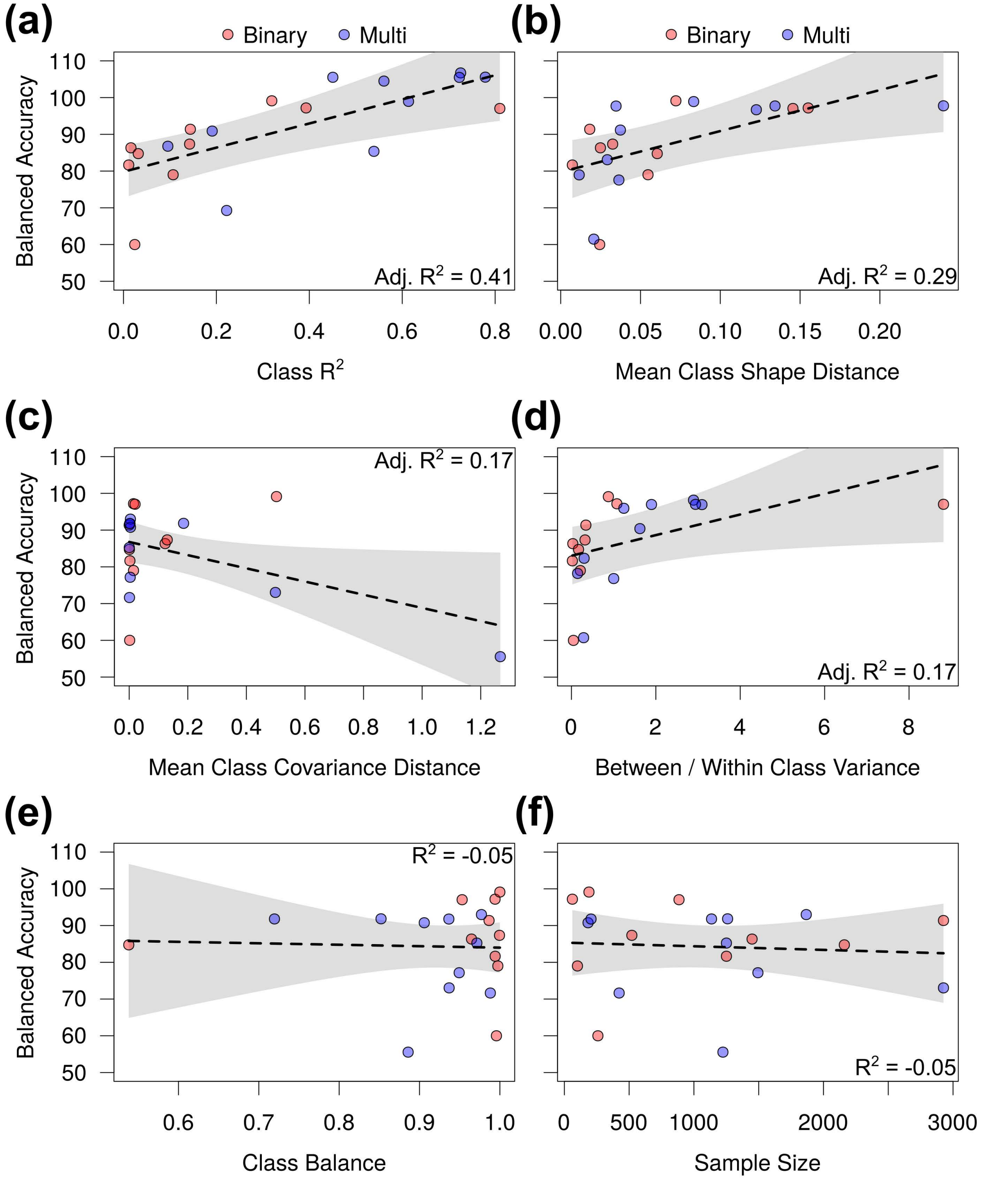


**Figure S6**. Balanced accuracy multiple regression plots that included a classification task covariate (red, binary; blue, multi) plus (a) class $R^2$, (b) mean class shape distance, (c) mean class covariance distance, (d) between- vs. within-class variance, (e) class balance, or (f) sample size explanatory variables. Lines of best fit with 95% confidence intervals are shown alongside model $R^2$ values.
